# Divergence in thermal performance contributes to ecotype maintenance in an intertidal snail: evidence from *in-situ* transplants

**DOI:** 10.64898/2026.07.21.739872

**Authors:** Christopher Dwane, Rodrigo Lorenzo, Juan Galindo, Emilio Rolán-Alvarez, Manuela Truebano

## Abstract

Physiological adaptation across environmental gradients can contribute to ecological speciation by limiting performance outside locally optimal habitats. Intertidal systems provide strong natural thermal gradients, yet the extent to which thermal physiology contributes to divergence across shore height remains poorly resolved. We investigated cardiac thermal performance in two ecotypes of the marine snail *Littorina saxatilis* occupying different shore heights along the Galician coast (NW Spain): a wave-adapted ecotype on the lower and mid- shore and a crab-resistant ecotype on the mid- and upper shore. Using infrared photoplethysmography, we quantified heart rate responses in both a reciprocal field transplant experiment and laboratory thermal ramping trials. In the field, the Wave ecotype exhibited significantly higher heart rates than Crab ecotype snails under native mid-shore conditions and after 1 day of exposure to upper-shore conditions. However, after 4 days of exposure to the upper shore, Wave ecotype snails showed a marked reduction in cardiac activity, whereas Crab ecotype populations maintained stable heart rates across transplant locations and durations. In laboratory ramping experiments, Crab ecotypes displayed lower baseline cardiac activity and greater thermal insensitivity across the rising phase of the thermal response curve, while the Wave ecotype exhibited higher cardiac performance and an earlier decline in heartrate at high temperatures. Together, these results demonstrate pronounced ecotype divergence in cardiac thermal physiology and suggest that chronic exposure to upper-shore conditions compromises cardiac performance in the Wave ecotype. Such physiological differences likely contribute to vertical zonation and the evolution of barriers to gene flow between these ecotypes.

## 1. Introduction

Physiological divergence in response to environmental thermal stress gradients is a key driver of the ecological diversification of life on Earth (Bartholomew, 1987; Chown, 2004; Spicer & Gaston, 1999). Because environmental stress selects for genotypes best suited to local conditions, populations of the same species may become physiologically specialised to local thermal conditions, restricting their ability to colonise other habitats and thereby acting as a barrier to gene flow (Kawecki & Ebert, 2004; Nosil, 2012). The potential role of physiological divergence in driving local adaptation and speciation has therefore long attracted the attention of ecophysiologists (Feder et al., 1987; Ghalambor, 2006; Janzen, 1967). Comparisons of related species living across stress gradients may be used to infer the role of physiological divergence in historic speciation events (Bartholomew, 1987; Keller & Seehausen, 2012), but such comparisons are often confounded by phylogenetic differences and species-specific evolutionary histories (Rezende & Diniz-Filho, 2012; Rundle & Nosil, 2005). By contrast, ecotype systems - where locally adapted populations of the same species exhibit genetic and phenotypic differences accompanied by partial but incomplete barriers to gene flow - offer an invaluable means to study the early stages of divergence and the formation of reproductive isolation in the face of gene flow (Johannesson et al., 2024; Rundle & Nosil, 2005). Although temperature adaptation has been implicated as a mechanism promoting population divergence in several ecotype systems (Gibbons et al., 2016; Kavanagh et al., 2010; Keller & Seehausen, 2012; Ohlberger et al., 2013), key challenges remain in understanding its role in the speciation process. In particular, although ecotypes sometimes display differences in thermal sensitivity, it is frequently unclear whether these differences actively contribute to maintaining divergence or instead represent correlated responses to other selective pressures (Keller & Seehausen, 2012). Furthermore, there are challenges in assessing the relative contribution of thermal selection in scenarios where divergent selection is occurring in response to multiple environmental factors (multidimensional divergent selection; White & Butlin, 2021). These uncertainties are compounded by the difficulty of linking laboratory measurements of thermal performance to the complex and dynamic stress regimes experienced in natural environments.

Intertidal environments are ideally suited to study local adaptation to thermal stress because they represent one of the most dramatic environmental clines in nature, characterised by a transition from terrestrial to fully marine conditions over a few vertical metres (Leeuwis & Gamperl, 2022; McMahon, 1990; Tomanek & Helmuth, 2002). Cardiac activity is a widely utilised proxy of thermal performance in intertidal hard-bodied organisms, including gastropods, as it underpins aerobic metabolic performance (Frederich & Pörtner, 2000), is highly responsive to temperature (Dong et al., 2021), and can be measured non-invasively (Burnett et al., 2013; Lima et al., 2025). Under increasing temperatures, heart rate in intertidal species typically follows a response similar to a classic thermal performance curve (TPC) (Dwane et al., 2023; Liao et al., 2021; Monaco et al., 2017), increasing with temperature to a peak (the thermal peak, T_peak_) before declining sharply at higher temperatures until cardiac flatline is reached. Organisms adapted to different shore heights may display both horizontal and vertical shifts in their cardiac TPCs. Species or populations occupying upper shore habitats are typically able to maintain cardiac function up to higher critical temperatures than those from lower shores, enabling them to resist greater thermal stress intensities and mobilise an effective heat-stress response (Dong et al., 2021; Stenseng et al., 2005; Stillman & Somero, 2000; Verberk et al., 2016). At the same time, in absolute terms, they may exhibit reduced heartrates across some or all of the TPC as a form of adaptive metabolic depression, enabling them to reduce metabolic energy expenditure at low tide (Hui et al., 2020; Liao et al., 2021; McMahon et al., 1995; Verberk et al., 2016; Wang et al., 2022). Such responses are especially important in upper shore habitats, where hard-shelled organisms with soft tissues may cease feeding and locomotion during low tide to limit water loss from desiccation, reducing energy intake (McMahon, 1990; Sokolova et al., 2000; Verberk et al., 2016). In contrast, lower shore organisms may be predicted to maintain comparatively higher heartrates at lower temperatures, reflecting elevated metabolic activity and facilitating more active lifestyles likely to be advantageous on the competition-dominated lower shore (Connell, 1961a, 1961b; McMahon, 1990; Monaco et al., 2017; Sokolova & Pörtner, 2001). Despite extensive physiological work characterising divergence in cardiac and metabolic activity in populations and species from different shore heights (Dong et al., 2021; Monaco et al., 2017; Sokolova & Pörtner, 2001; Stenseng et al., 2005; Stillman & Somero, 2000; Tomanek & Somero, 2000), how these physiological differences translate to performance under natural, fluctuating field conditions remains poorly resolved.

A well-studied example of intertidal vertical zonation within a single species is found in populations of the rough periwinkle *Littorina saxatilis* in Galicia (NW Spain). Here, two ecotypes are present at different tidal levels in response to divergent selection: a larger Crab ecotype characterised by a robust, crab-resistant shell, and a smaller Wave ecotype, characterised by a smooth, thin shell and high foot tenacity conferring resistance to wave action (Rolán-Alvarez, 2007). Vertical zonation and genetic divergence in the two ecotypes are thought to be maintained primarily by differences in the selective forces of wave action and predation present at the two shore heights, with the Wave ecotype being associated with a mussel belt on the lower shore and the Crab ecotype with the barnacle zone on the upper shore (Butlin et al., 2014a). However, the presence of a hybrid zone in the mid-shore, where both ecotypes coexist among patches of mussels and barnacles, complicates this scenario by allowing for heterospecific matings and gene flow. Both ecotypes remain distinct within this hybrid zone, although recent evidence suggests that Crab ecotype snails that live close to the hybrid zone are genetically and phenotypically distinct from those of the upper shore, suggesting some introgression and the potential for adaptive within-ecotype differences across shore height (Raffini et al., 2025). Recently, renewed focus has been placed on whether differences in physiology of the two ecotypes may play an important role in their vertical zonation and genetic differentiation, in addition to predation and wave action (Dwane et al., 2021; Johannesson et al., 2024; Morales et al., 2019). Recent work has established that the Wave ecotype possesses increased sensitivity to temperature change, and reduced tolerance of extreme temperatures, compared to Crab ecotype snails from the upper shore, suggesting that thermal stress physiology may play a hitherto underexplored role in maintaining the selective pressures leading to the maintenance of divergence between the two ecotypes (Dwane et al., 2021). However, it remains unclear how these physiological differences manifest across shore height under natural conditions, where multiple stressors such as heating rate, exposure duration, and desiccation may compound physiological stress in ways that are difficult to predict from single-stressor laboratory experiments (Marshall et al., 2010; McMahon, 1990; Tagliarolo & McQuaid, 2016).

In this study, we aimed to bridge the gap between our understanding of thermal physiology in the two Galician *L. saxatilis* ecotypes, and their ecological divergence across shore height, by examining cardiac thermal responses under both field and laboratory conditions. We conducted a reciprocal transplant experiment using snails from the upper shore (Crab ecotype) and mid-shore hybrid zone (Crab and Wave ecotypes) over multiple days during thermally extreme summer low tides, measuring *in-situ* heart rate responses at short (1 day) and longer (4 day) exposure periods. To complement this field experiment, we performed a laboratory experiment to characterise cardiac thermal performance curves across ecotypes and populations during an acute thermal ramp. By including Crab ecotype populations from both shore heights, we were able to distinguish effects of potential fine-scale adaptation across shore height (upper shore versus mid-shore) from those associated with ecotype identity (Crab versus Wave). Specifically, we aimed to test the prediction that both mid- and upper shore Crab ecotype snails would exhibit consistent physiological responses across shore heights, whereas the mid-shore Wave ecotype would display greater thermal sensitivity of heart rate, reduced thermal limits, and signs of physiological stress following multi-day exposure to upper-shore conditions.

## 2. Methodology

### 2.1. Field Experiment

#### 2.1.1. Study Site Selection

The *in-situ* reciprocal transplant experiment was conducted at Centinela on the west coast of Galicia, Spain (N 42°04’38.06” W 8°53’47.47”)(Raffini et al., 2025). The experiment was carried out during a 9- day window in July 2022, in which low spring tides occurred in the morning or early afternoon, consequently ensuring high temperatures were experienced during emersion.

Three replicate sites spaced horizontally at approximately 30 m intervals were selected at each shore level (mid and upper shore; Fig. 1A). Mid-shore locations were defined based on the presence of both Wave and Crab ecotypes (Fig. 1B), thus representing the hybrid zone identified by previous studies (Raffini et al., 2025); this habitat was characterised by a mosaic of barnacle and mussel patches (Carballo et al., 2005). Upper shore locations were positioned inland and at higher elevation on the shore and defined based on the presence of only Crab ecotype snails; this habitat consisted of barnacle patches only (Fig. 1A-B). To reduce the impact of predation on sample size, upper site locations were chosen in areas away from larger rock overhangs inhabited by the predatory crab *Pachygrapsus marmoratus*. Temperature data loggers (ElectricBlue, Vairão, Portugal) were placed at each site (n = 5 on the upper shore, 5 on the lower shore; 1-2 per site) to continuously record field temperatures (± 0.1°C; 10 min frequency) during the experimental period.

**Figure 1:**
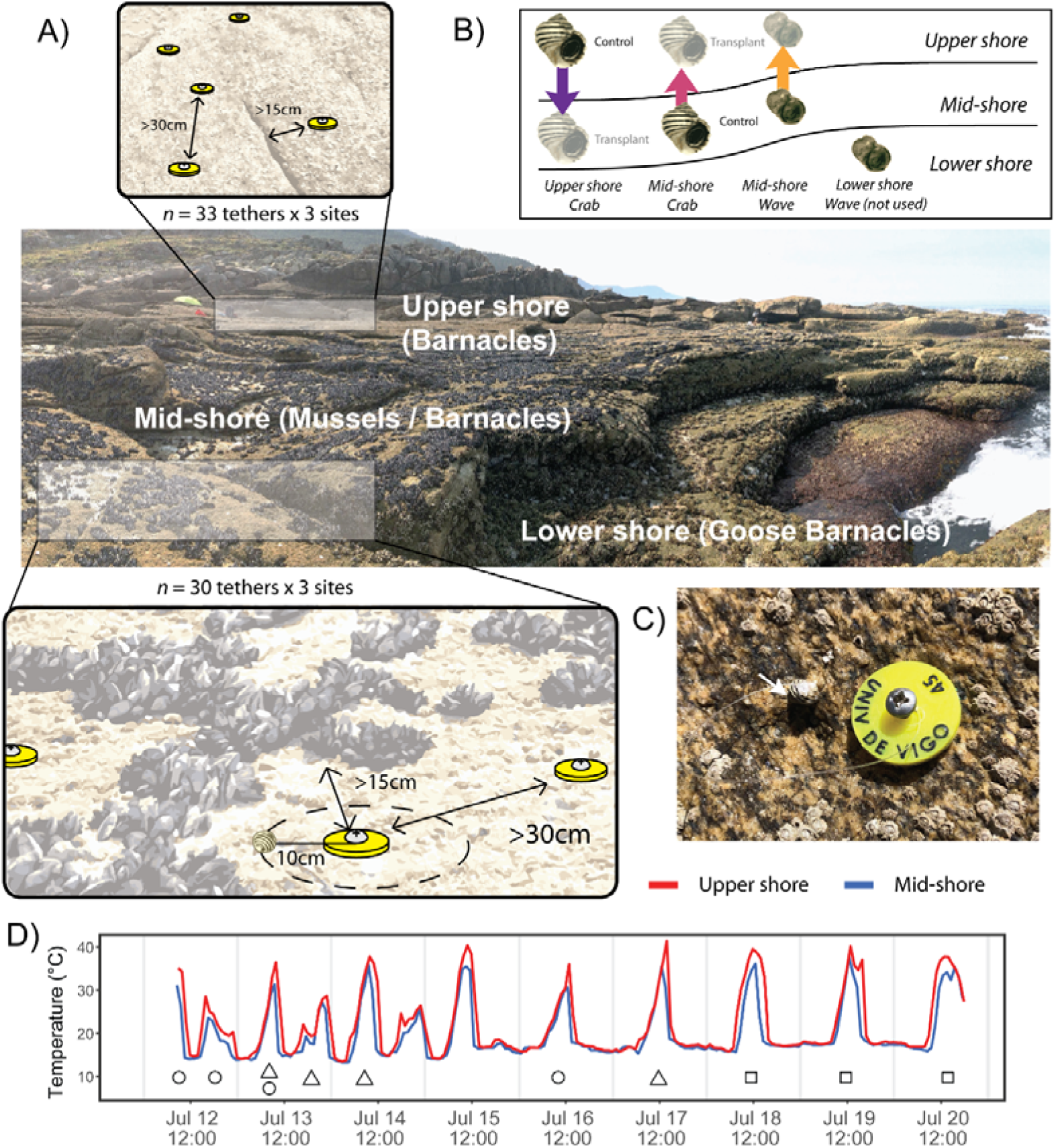
Experimental design of the reciprocal transplant. (A) Schematic showing the vertical position of upper and mid-shore locations used in the experiment, which were defined based on habitat type. Note that for each shore height, three replicate sites were chosen, spaced approximately 30m apart along the shoreline. Tether attachment sites on the mid-shore can be seen in the bottom left corner. Inset figures depict placement of tether attachment locations on the upper and mid-shores, showing minimum spacing between tethers (>30 cm). At both shore heights, attachment locations were placed in barnacle-dominated substrate, positioned at least 15cm distant from either crevices or mussel beds on the upper and mid-shores respectively. Snails were attached to 10cm long tethers, preventing them from moving outside the barnacle habitat. Inset figures are not to scale. (B) Conceptual diagram of reciprocal transplants across shore heights, distinguishing control and transplant treatments for each ecotype/ shore height combination. (C) Photograph of an individual experimental snail from the Upper shore Crab population (indicated by white arrow) attached to its tether (monofilament fishing wire) on the upper shore. (D) Averaged temperature profiles for the upper shore (n= 5 loggers) and mid-shore (n=5 loggers). Data is downsampled to an hourly frequency to improve visual presentation. Shapes indicate deployment and sampling dates for the three experimental batches (circle = batch 1; triangle = batch 2; square = batch 3). From left to right, the shapes for each batch indicate the timepoints at which: 1) snails were collected from the field; 2) snails were returned to the shore and placed at reciprocal transplant locations; 3) heartbeat recordings were taken for the acute (1-day) exposure period; 4) heartbeat recordings were taken for the chronic (4-day) exposure period. Note that for batch 3, recordings were taken for the acute exposure period only.

At each site, attachment points for snail tethers (n = 30 - 33 per each of six sites, three per shore level) were created by drilling holes into the rockface and inserting screws spaced approximately 30 cm apart in flat, barnacle-dominated areas, at least 15 cm away from crevices and mussel patches (Fig. 1A, insets). The latter were avoided to minimise refuge seeking and behavioural thermoregulation. Although the mid-shore is a mosaic of mussel beds and barnacles, the Wave ecotype is found across both habitat types (Carballo et al., 2005). Therefore, the use of barnacle patches for attachment points across both shore heights offered appropriate habitat for both ecotypes. Screws were marked with numbered acrylic tags for identification (Fig. 1C).

#### 2.1.2. Snail Collection and Transplantation

The transplantation experiment was performed using three separate batches of snails starting on different days (batches 1, 2, and 3; Fig. 1D, see accompanying caption). As the sampling procedure for collecting heart rate data was-time-intensive and limited by tidal windows, conducting the experiment across multiple batches enabled the collection of data from more individuals. For each batch, snails from the three populations—mid-shore Crab, mid-shore Wave, and upper-shore Crab—were collected at low tide from their respective shore heights on either the 12^th^, 13^th^ or 18^th^ July (Fig. 1D). Note that throughout, we use “population” to refer to the combination of ecotype (Wave or Crab) and the shore height it was collected from - thus, “mid-shore Crab” represents the same ecotype as “upper shore Crab”, but from a different shore height (Fig. 1B). Immediately following collection, snails were transported to the Toralla Marine Science Station (ECIMAT). Each was then fitted with a 10 cm tether made of fishing wire attached to the shell with superglue (Loctite) and held in a flow through seawater system at ambient seawater temperature. On the subsequent daytime low tide, snails were returned to the shore for reciprocal transplantation. In the case of batches 1 and 2, snails were transplanted at evening low tide on the same day they were collected (Fig. 1D, see accompanying caption). For batch 3, the next low tide occurred at night; thus, snails were instead transplanted at low tide the following day (Fig. 1D, see accompanying caption). To perform the transplantation, snails from each subpopulation were tethered to screws at each of the three upper and three mid-shore sites (Fig. 1C). In total, 309 snails were transplanted across the three batches (mid-shore Crab, n= 102; mid-shore Wave, n= 103, upper shore Crab, n=104). This number included numerous additional individuals not used for subsequent heart rate data collection, to account for anticipated losses due to mortality, predation or broken tethers. Note also that individual screw locations were in some cases used multiple times throughout the three batches.

#### 2.1.3. Field Measurements

To measure thermal performance under acute (single day) vs. chronic (multi day) exposure to transplant conditions, heartbeat recordings were obtained from tethered snails at each site during low tide one day post-transplantation for all three batches (acute exposure), and again four days post-transplantation for batches 1 and 2 (chronic exposure; Fig. 1D). On each day in which recordings were taken, infrared sensors (CNY70, Vishay, USA) were attached to six snails from each site (two from each population per site) and held in place using a small amount of dental wax placed over the sensor tip. Immediately prior to heartbeat recordings, a thermal image of the snail attached to the sensor was captured using a thermal imaging camera (Testo 883, Testo SE & Co. KGaA, Germany) at 95% emissivity and stored to generate reference body temperatures for each heartbeat recording (Seuront et al., 2018). Heartbeat recordings were then captured over a 2-minute period at 40 Hz using a system consisting of the infrared sensor attached to a heart rate amplifier (AMP03, Newshift, Portugal) connected to a data acquisition device (USB-6001, National Instruments, USA) and laptop. To shield the sensors from solar infrared radiation during readings, small metal cups lined with aluminum foil were briefly placed over the sensors and animals during heartbeat recordings and were then removed immediately afterwards.

As temperatures increased over the course of emersion at low tide, two heartbeat recordings (2 min each) were captured from each snail at different times during the emersion period to capture performance across a range of temperatures *in-situ*. The amount of time between the two recordings varied due to setup time involved in taking readings between different locations, but was at least 30 min (average time between recordings: 152 ± 66 min; mean ± sd). Note that this means that each snail was sampled twice and these repeated measures were used in the statistical analysis. Sensors were left attached to the snails between heartbeat recordings. After the last measurement had been taken on day one, sensors and dental wax were detached from the snails, which were left tethered on the shore. In total, recordings were taken from 154 snails. Where possible, the same snails which had been used for measurements on day one were used again for day four; however, losses of tethered snails over the four-day period were high and as a result only a total of 19 snails out of 102 measured on day one in groups one and two were reused on day four.

Therefore, the majority of heart rate recordings on day four were instead taken from additional tethered snails which had not been used on day one. Losses were likely due to a combination of predation (especially on the high shore) and mechanical detachment of the snail from the tethers due to wave action, but as these effects could not be readily distinguished, data on the relative causes of losses could not be obtained.

Reference body temperatures for each heartbeat recording were obtained from the thermal image captured immediately prior to recording and analysed using the IRSoft software (TESTO, Testo SE & Co. KGaA, Germany). For each image, a polygon or oval was manually drawn around the visible body and shell of the animal, and the mean surface temperature of all pixels within this region of interest was calculated. This value was used as an estimate of individual body temperature in subsequent analyses.

### 2.2. Laboratory Experiment

To complement field heartbeat recordings and assess differences in cardiac activity between populations in response to acute warming, a laboratory experiment was conducted using the facilities at ECIMAT (Toralla Marine Station, University of Vigo, Vigo, Spain). Snails from each of the three populations were collected from their respective field locations and transported to ECIMAT, where they were held for ∼24 h in glass aquaria (18 x 10 x 6 cm) connected to a flow-through seawater system at ambient seawater temperature (temperature = ∼20°C). The following day, snails were fitted with infrared sensors (CNY70, Vishay, USA) as previously described and placed in dry falcon tubes partially submerged within a static water bath (FisherBrand Isotemp, Fisher Scientific, UK) at 20 °C. Following a 30 min period to allow snails to settle into the tubes and remove handling stress, the heartbeat of each individual snail was recorded for 1 min using the same infrared logging system used in the field experiment. As readings could only be taken from six snails at a time (corresponding to the number of channels available on the USB-6001 data acquisition device) recordings were taken sequentially from batches of six snails (two from each population) until measurements had been obtained from all individuals. After recording all individuals at 20 °C, the setpoint on the bath was raised by 2 °C, to which bath temperatures would equilibrate within 5 min. Once the desired temperature was reached, animals were held at this temperature for 10 min, after which heartbeat recordings were obtained as described from each batch of six individuals in sequence. This process was repeated at 2°C increments every 30 min until a temperature of 40 °C was reached, following which the experiment was ended. The thermal ramping was performed in two runs on different days, with n = 24 and n = 36 snails respectively, divided evenly across the three populations.

Because of the thermal properties of the bath, falcon tubes and the surrounding air, the temperatures at which the falcon tubes equilibrated at each setpoint varied by ± 0.3°C (in one instance ± 0.7°C) from the set temperature of the bath. Accordingly, the actual temperature experienced by the snails at each setpoint (measured using an HH806U temperature logger; Omega, USA) was used in preference over the setpoint temperature in data analysis.

### 2.3. Data Processing

For both the field and laboratory experiment, heart rates were obtained from raw voltage signals using a custom-built R shiny application which allowed semi-automated identification of individual heartbeats within a user interface. Each reading was first smoothed using the “ksmooth” function, following which individual peaks were identified using the “findpeaks” function. The application automatically adjusted the bandwidth of the smoother (between 0.1 and 0.6) to result in the fit that gave the lowest standard deviation of heart rate frequency (in Hz). This range of bandwidths matches recommendations from published r-based approaches for detecting peaks from heart rate data in intertidal species (Lima et al., 2025).

In the vast majority of cases this resulted in an optimal fit of the smoother to the raw data based on manual observation; however, in instances where the fit was visually poor, the bandwidth could be manually adjusted. Signals containing minor noise were prefiltered using a Bartlett window set to 20 samples. The fit of the calculated heartbeats to the raw data was visually inspected to assess the quality of the heart rate data. Recordings containing poor quality, highly irregular, or missing heartbeat signals were discarded from further analysis. In total, of 336 recordings taken during the field experiment, 45 were discarded for the aforementioned reasons, while for the ramping experiment in the laboratory, data from eight individuals out of 60 were discarded due to poor or unreadable signals across some or all of the temperature range.

### 2.4. Data Analysis

#### 2.4.1. Field Experiment

All statistical analysis was performed in R and Rstudio. To compare cardiac activity under one day vs. four-day reciprocal transplantation across subpopulations and shore heights, a linear mixed-effects model was used, with heart rate (bpm) as the response variable and population (mid-shore Crab, mid-shore Wave, upper shore Crab), shore height (mid vs. upper) and transplant duration (one day vs. four day) as categorical predictor variables, and body temperature as a linear covariate. An interactive effect of temperature was considered in the initial model but was removed during model simplification. For random effects, snail identity (to account for the fact that multiple readings were taken from individuals both within and between duration timepoints), Batch (denoting the day on which the snails were deployed in the field: 1, 2, or 3, as outlined in Section 2.1.2.) and replicate site (three per shore height) were included. The latter two were included to account for shared environmental conditions experienced by batches and potential non-independence among individuals within sites. Size (measured from spire to aperture tip, in mm) was included as a covariate in the initial model, but was removed from the final model based on model selection. Model suitability was evaluated using posterior predictive tests in the “Performance” package (Lüdecke et al., 2021) and diagnostic plotting of simulated residuals using “DHARMa” (Hartig, 2023), with model simplification performed based on AIC values. The final model was constructed using the “lmer” function in the package lmerTest (Kuznetsova et al., 2017) and tested using ANOVA with type III sum-of-squares. Post-hoc testing was performed in the package “emmeans” (Lenth & Piaskowski, 2025), with planned contrasts comparing differences between ecotype and transplant duration within each shore height.

To examine effects of experimental exposure duration on thermal sensitivity of heart rate across ecotypes and transplant conditions, Q_10_ values were calculated using the repeated heart rate recordings made at different temperatures from each snail on a given day, using the equation:

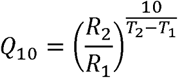

With R1 and R2 being the heart rate values and T1 and T2 being the temperatures during the first and second recordings, respectively. Because Q_10_ values are extremely sensitive to fluctuations or errors in rate measurements when calculated over small temperature increments, we only calculated Q_10_ for individuals where a difference in temperature of over 4 °C was observed between pairs of recordings, resulting in a dataset where the median temperature difference between the start and end temperatures was 7.9 °C (interquartile range: 6.2 - 9.9 °C). Q_10_ values were compared statistically across treatment groups using a linear model with population, shore height and transplant duration as predictor variables. Random effects of Size, Individual, Group and Site were initially included in the model but were subsequently removed as they did not improve model fits to data. In addition, because the sample size of the final dataset for field Q_10_ (n= 92 samples across 12 groups) was quite small relative to the original dataset due to the restrictive sampling criteria used, a post hoc sensitivity analysis was conducted to estimate the statistical power of the model to detect the Ecotype × Height × day interaction. This was performed using the pwr.f2.test function in the R package “pwr”.

#### 2.4.2. Laboratory Experiment

Initial visual inspection of the data revealed that while cardiac responses for the mid-shore Wave ecotype displayed a characteristic TPC-shaped response, performance for the upper shore Crab and mid-shore Crab ecotype was still increasing when the experiment was terminated at 40°C, meaning they did not display a thermal optimum within the range of temperatures measured. Because fitting full thermal performance curves was therefore not appropriate for all populations, we instead analyzed temperature-dependent variation in heart rate using Generalized Additive Models (GAMs). To evaluate whether variation in cardiac thermal performance was best explained by ecotype differentiation, shore height, or population-specific differences incorporating both factors, we compared three alternative GAMs using Akaike’s Information Criterion (AIC). The first (Model 1) included a unique smoother for each population (mid-shore Crab, mid-shore Wave and upper shore Crab) to allow the relationship between heart rate and temperature to differ across both ecotypes and shore heights. The second model (Model 2) fitted separate smoothing terms for each ecotype (Crab vs. Wave), while the third (Model 3) fitted separate smoothing terms based on shore height (upper vs. mid-shore). Each model included snail identity as a random effect with a random intercept to account for individual differences in heart rate, while a fixed effect of population was included to allow for differences in intercepts between the three populations. A covariate effect of shell length was incorporated to account for the effect of potential within-population in size. GAMs were fitted using the mcgv package, while model comparisons were performed using the “anova” and “AIC” functions in base R.

In addition, to compare thermal sensitivity of heart rate across the three populations and provide a direct comparison with the results from field data, we calculated Q_10_ values for snails used in the ramping experiment following the protocol described in section 2.4.1., but instead using heart rate values recorded at temperatures corresponding to setpoints of 20°C and 30°C. This 10°C temperature range approximately matches the median start and end temperatures used to calculate used to estimate Q_10_ values from the field dataset (20.85 and 29.80°C, respectively), and also while still encompassing the rising slope of the TPC across all three populations based on visual inspection of the data (see Results). A linear model with Post-hoc Sidak tests was used to compare Q_10_ values between the three populations. Statistical power of the model was assessed using the function “pwr.anova.test” in the R package “pwr”.

## 3. Results

### 3.1. Field Experiment

The linear mixed-effects model comparison of cardiac performance across transplant conditions revealed a significant three-way interaction between Population x Shore height x Transplant duration (χ² = 6.312, df = 2, p = 0.043), and a significant interaction between Population x Transplant duration (χ² = 13.585, df = 2, p < 0.001), on cardiac activity. A significant main effect of Population (χ² = 70.542, df = 2, p < 0.001) and covariate effect of temperature (χ² = 56.465, df = 1, p < 0.001; Table S1), were also detected, while the effect of shore height alone was not significant.

Heart rate increased with temperature across all groups (Fig. 2), but clear differences were observed between ecotypes. Following acute (1 day) transplantation to either shore height, mid-shore Wave ecotype snails exhibited significantly higher heart rates than both Crab ecotypes. (Fig. 2; Table S2). However, after 4 days of exposure to conditions on the upper shore, mid-shore Wave ecotype snails showed a pronounced reduction in heart rate (estimated marginal mean: 73.315 ± 5.103 SE) compared to day 1 (98.473 ± 4.709 SE), converging with values observed in Crab ecotypes. In contrast, neither mid-shore nor upper-shore Crab ecotypes exhibited significant changes in heart rate between 1- and 4-day exposures, and Wave ecotype snails maintained on the mid-shore also showed no change over time (all post-hoc Sidak comparisons, p > 0.05; Fig. 2; Table S2).

**Figure 2:**
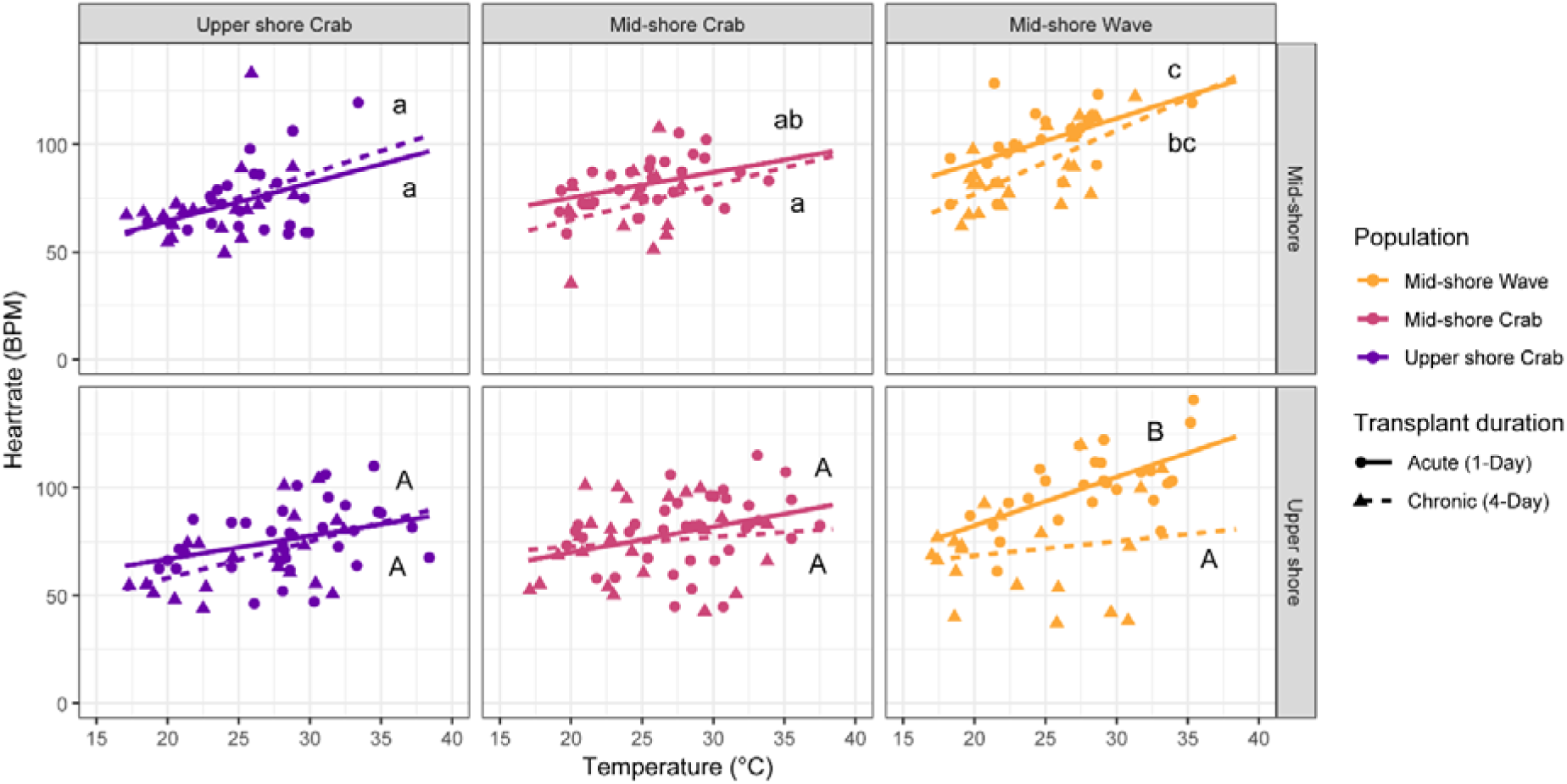
Comparison of field heart rate measurements from *L. saxatilis* snails used in the reciprocal transplant experiment, plotted against the temperatures at which they were recorded. The top row of panels indicates measurements from snails exposed to mid-shore conditions during reciprocal transplant, while the bottom panels display measurements from snails held exposed to the upper shore. Groups sharing the same letter did not differ significantly from one another (p > 0.05, based on post-hoc tests). Note that post-hoc tests were performed across populations and transplant durations within (not between) shore heights, as shown by the use of upper- and lower-case letters for the upper and mid-shore, respectively.

Thermal sensitivity (Q_10_) of cardiac activity measured in the field did not differ significantly among populations, transplant durations, or shore heights, and no interactive effects were detected (all p values > 0.05).Visual comparison of Q_10_ values suggested a pattern similar to that observed in the field heart rate data, with mid-shore Wave ecotype snails displaying reduced Q_10_ following chronic transplantation on the upper shore (Fig. 3). However, the population × transplant duration × shore height interaction had a small effect size (partial η² = 0.03; Cohen’s *f* = 0.18), and post-hoc sensitivity analysis indicated that the sample size (n = 92) provided approximately 27% power to detect an effect of this magnitude. This suggests that population- and environment-dependent effects on thermal sensitivity are relatively small, and that much larger sample sizes would be required to reliably detect any effects.

**Figure 3:**
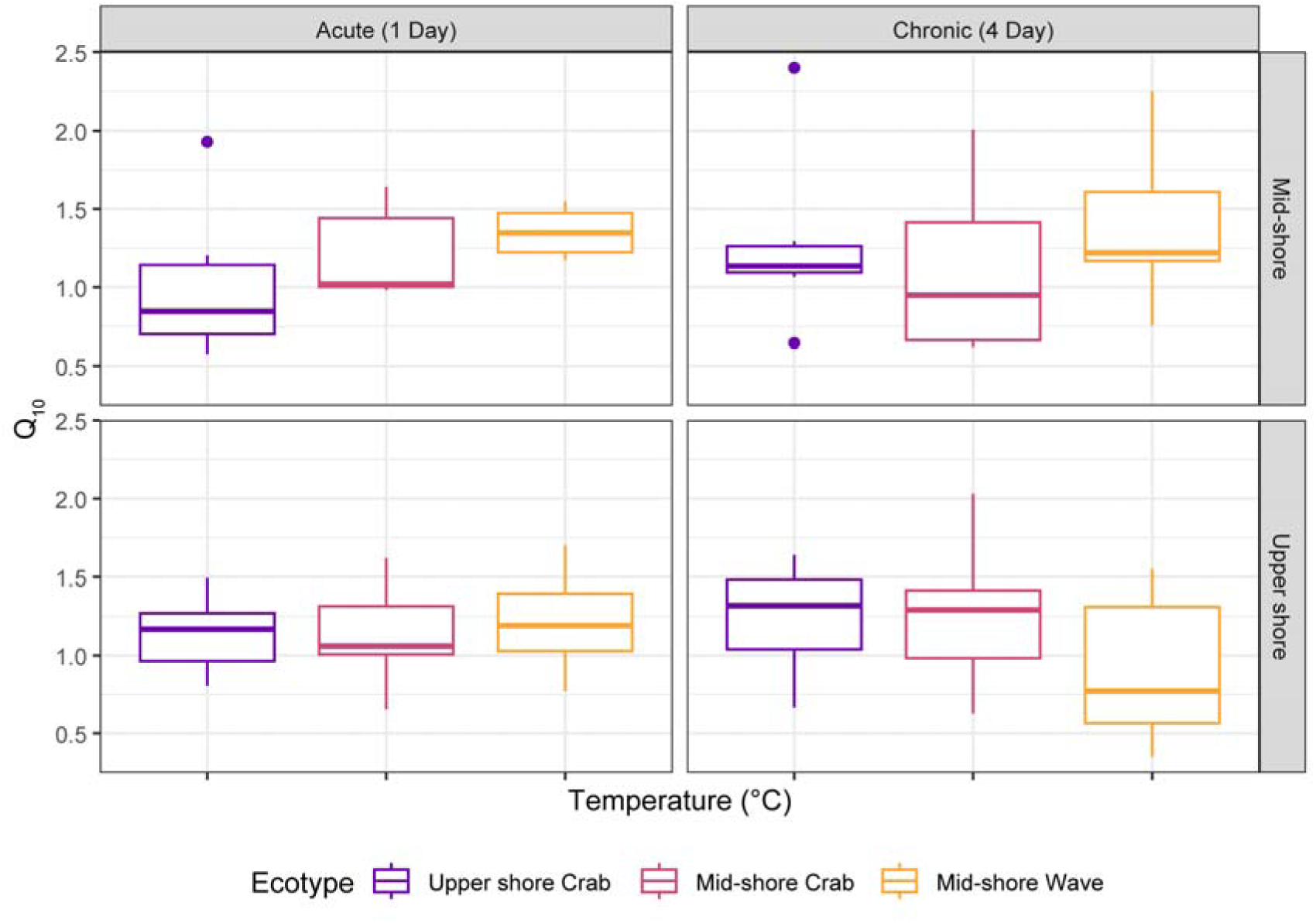
Thermal sensitivity of cardiac activity (Q_10_) in *L. saxatilis* across shore height, population, and transplant duration in the field experiment. Q_10_ values were calculated for individual snails using paired heart rate measurements obtained during natural heating over emersion, where the temperature difference between recordings exceeded 4 °C (median difference: 7.9 °C). Data are shown for three populations (mid-shore Wave, mid-shore Crab, upper-shore Crab) across two shore heights (mid vs. upper) and two exposure durations (1 day and 4 days post-transplant). Sample sizes are as follows: **upper shore Crab:** acute, mid-shore, n = 6; chronic, mid-shore, n=7; acute, upper shore, n = 10; chronic, upper shore, n=8; **mid-shore Crab:** acute, mid-shore, n = 7; chronic, mid-shore, n=4; acute, upper shore, n = 12; chronic, upper shore, n=9; **mid-shore Wave:** acute, mid-shore, n = 6; chronic, mid-shore, n=9; acute, upper shore, n = 6; chronic, upper shore, n=8.

### 3.2. Laboratory Experiment

Comparison of candidate GAMs indicated that variation in cardiac performance was best explained by differences between ecotypes rather than shore height of origin. Model fits did not differ significantly between Model 1, in which separate smoothers were fitted for each of the three populations, (mid-shore Crab, mid-shore Wave, upper shore Crab), and Model 2, in which separate smoothers were fitted by ecotype (wave vs. Crab) (Model 1 AIC = 4633.070; Model 2 AIC = 4633.712; F = 1.68 p= 0.17). By contrast, Model 3, in which smooth terms varied by shore height (upper vs. mid-shore), performed significantly worse than both (AIC = 4703.862; Model 1 vs Model 3: F = 15.46, p < 0.001; Model 2 vs Model 3: F = 9.984, p < 0.001). This indicates that differences in the relationship between heart rate and temperature are primarily associated with ecotype identity rather than shore height.

Across the thermal ramp, heart rate increased with temperature in all groups, but the shape and magnitude of responses differed between ecotypes (Fig. 4). Wave ecotype snails exhibited higher baseline heart rates and a distinct thermal response, reaching a peak within the experimental temperature range. In contrast, both Crab ecotypes showed lower cardiac activity and a more gradual increase in heart rate with temperature, with no clear peak observed up to 40°C. Post-Hoc comparison between populations using the best supported model (Model 1) revealed significantly higher intercepts in the Wave ecotype (estimated marginal mean: 93.82 ± 4.80 SE) compared to the upper- and mid- shore Crab ecotypes (48.76 ± 3.89 and 57.60 ± 3.5 respectively; p values <0.001), while the two Crab populations did not differ (p = 0.175), suggesting higher baseline heart rate in Wave ecotype snails.

**Figure 4:**
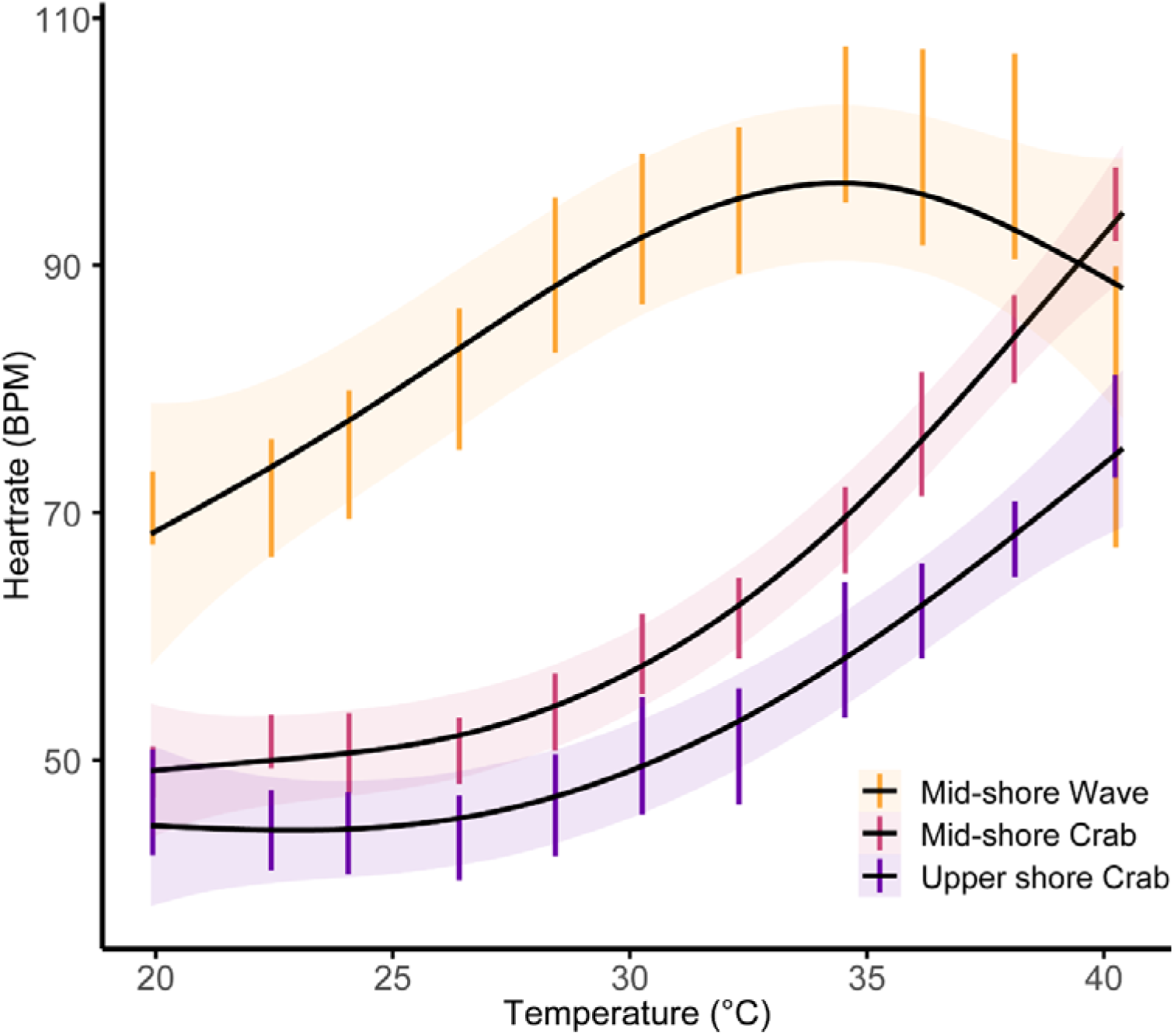
Cardiac performance in response to a thermal ramp across three populations of *L. saxatilis*. Points and bars indicate mean heart rate (BPM) ± SE for each temperature setpoint at which measurements were taken. Lines and shaded areas indicate Generalised Additive Model (GAM) fits ± SE, fitted using the ggplot2 package in R. Note that in this plot, a separate smoother was fitted for each population – thus, the plotted curves are analogous to Model 1 as described in Section 2.4.2. Results from Model 1 and Model 2 (in which the same smoother was applied for both Crab ecotype populations) did not differ significantly; see Section 3.2.

Consistent with field observations, Q_10_ values did not differ significantly among populations (F_2,48_ = 2.7404, p = 0.0749, Fig. 5). However, there was a moderate effect size of population (η² = 0.10; Cohen’s *f* = 0.34) and post-hoc sensitivity testing revealed an estimated power to detect a significant effect of 56% based on the sample size used, suggesting the potential for real differences in Q_10_ between populations that may have been undetected due to sample size limitations.

**Figure 5:**
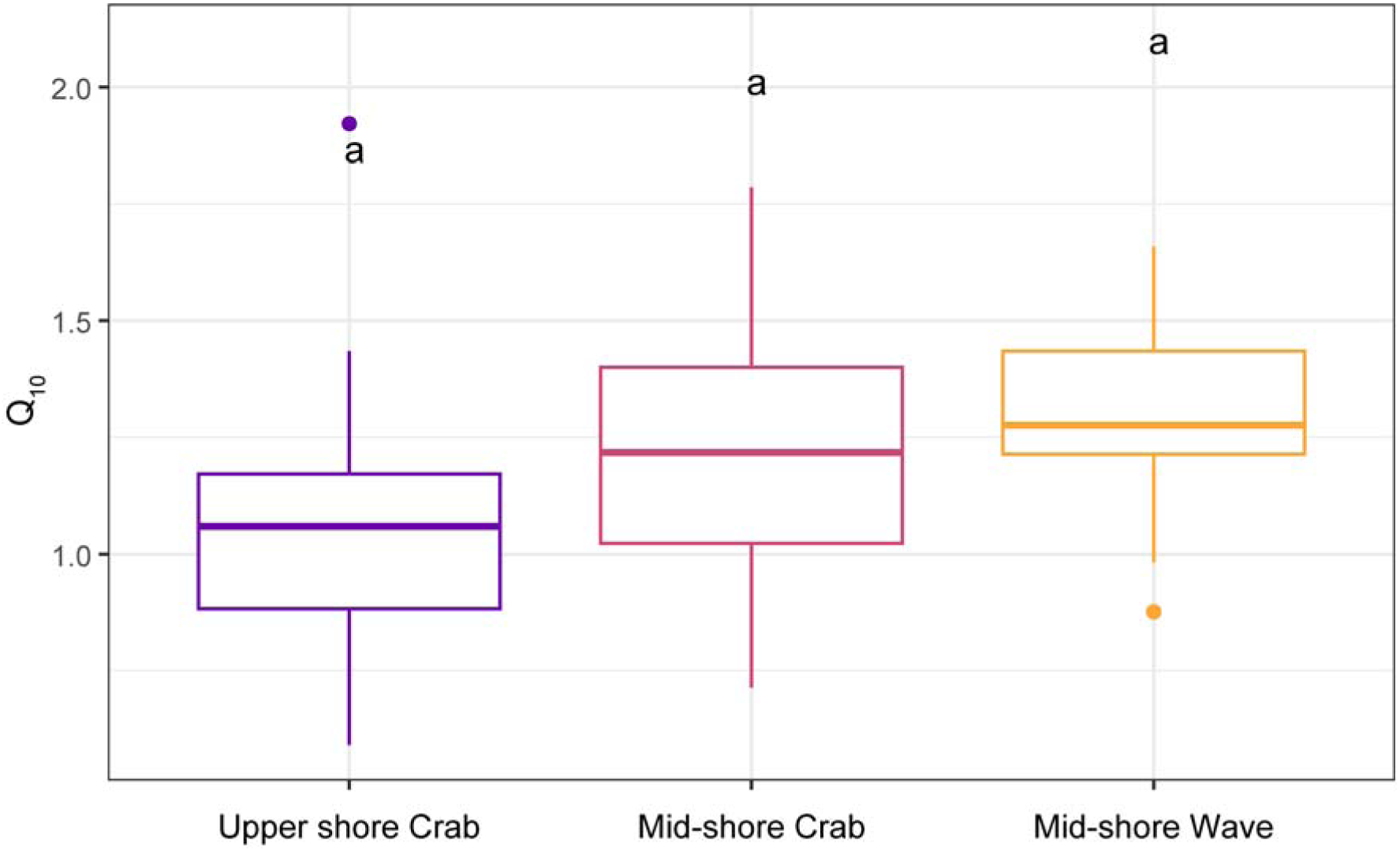
Thermal sensitivity of cardiac activity (Q_10_) in *Littorina saxatilis* populations measured during the laboratory thermal ramping experiment. Q_10_ values were calculated for individual snails using heart rate measurements obtained at 20 °C and 30 °C, corresponding to the rising phase of the thermal response curve across all populations. Data are shown for the three populations (**upper-shore Crab,** n= 18; **mid-shore Crab,** n= 18; **mid-shore Wave,** n = 16). Letters indicate significant differences at P < 0.05.

## 4. Discussion

The Galician *Littorina saxatilis* hybrid zone represents a striking pattern of divergence associated with environmental variation across shore height (Rolán-Alvarez, 2007). The parallel evolution of this and other hybrid zones across the species’ range (e.g. the Swedish coast) makes *L. saxatilis* a model system for studying incipient speciation and the formation of barriers to gene flow (Butlin et al., 2014b; Johannesson et al., 2024; Raffini et al., 2025). However, the physiological dimension of such divergence remains comparatively underexplored. In our reciprocal transplant experiment, Wave ecotype snails consistently exhibited higher heart rates than Crab ecotypes under native mid-shore conditions and after short-term (1 day) exposure to the upper shore, indicating intrinsic differences in physiological performance between ecotypes. However, after chronic (4 days) exposure to upper-shore conditions, Wave ecotype snails exhibited a marked reduction in heart rate, while no comparable changes were observed in mid-shore Crab ecotype snails transferred to the upper shore. This reduction in cardiac activity is consistent with a decline in physiological performance under prolonged exposure to upper-shore conditions, potentially reflecting a stress response or reduced condition arising from the combined effects of temperature, desiccation, and emersion. In addition to well-documented morphological and genetic differentiation between the Wave and Crab ecotypes (Galindo et al., 2013; Raffini et al., 2025), we suggest that divergence in cardiac thermal physiology, as a proxy for metabolic performance, constitutes an important component of local adaptation across this gradient, which may have physiological and ecological consequences for the maintenance of vertical zonation and morphological and genetic differentiation in the ecotypes.

Divergence in metabolic rate across shore height is a characteristic feature associated with the vertical zonation of intertidal species and populations (Dong et al., 2021; Leeuwis & Gamperl, 2022). In gastropods, metabolic depression is frequently observed in species inhabiting thermally stressful and unpredictable environments and may be reflected in reduced cardiac activity and thermal insensitivity of heart rate across a portion of the rising phase of the thermal performance curve (Marshall & McQuaid, 2011; Verberk et al., 2016). Alternatively, metabolic depression may manifest as a non-adaptive response to low energy reserves or poor condition (Leeuwis & Gamperl, 2022; Verberk et al., 2016). In our laboratory experiment, Crab ecotype snails exhibited lower cardiac performance, with no clear thermal optimum reached within the experimental range (Fig. 4). This pattern may be consistent with an adaptive reduction in metabolic activity associated with the more thermally stressful and variable upper shore environment. This aligns with comparisons of the performance of metabolic activity in high- and low-shore populations of *L. saxatilis* (Sokolova et al., 2000; Sokolova & Pörtner, 2001), as well as a latitudinal comparison incorporating this population which suggested that reduced cardiac performance in this population may be an adaptive response to greater thermal stress at lower latitude (Dwane et al., 2023). By contrast, the Wave ecotype exhibited higher cardiac activity and a distinct thermal response which peaked within the experimental temperature range. This is consistent with previous findings showing reduced critical thermal limits and greater thermal sensitivity in this ecotype (Dwane et al., 2021). In the present study, thermal sensitivity of cardiac activity (Q_10_) did not differ among populations in either field or laboratory experiments, potentially suggesting that differences among ecotypes may be more strongly associated with variation in absolute cardiac performance than with temperature dependence of heart rate. However, we cannot exclude the possibility that underlying differences in thermal sensitivity exist, especially as observed effect sizes for population in the laboratory experiment was moderate (η² = 0.10), suggesting that biologically meaningful differences in Q_10_ between populations may exist but were not detected due to lack of statistical power.

It is important to note that heart rate is an indirect proxy for metabolic rate, and size differences between the ecotypes and populations may account for some of the observed differences in cardiac performance we observed. However, biochemical and life-history evidence points to a requirement for the Wave ecotype to maintain higher metabolic activity levels under normal field conditions. Lower shore species often exhibit higher maximum activities of key metabolic enzymes (Sokolova & Pörtner, 2001) and other performance traits (Monaco et al., 2017), consistent with selection for increased feeding and activity in more energy-rich, competitive environments. The Wave ecotype is also morphologically adapted to resist high wave action on the lower and mid shore, including a foot muscle 1.4 times larger than the Crab ecotype (Martínez-Fernández et al., 2008). Proteomic comparison of the two ecotypes has demonstrated higher constitutive upregulation in the Wave ecotype of arginine kinase, an enzyme catalyzing the rapid regeneration of ADP into ATP during periods of intense activity, and fructose-bisphosphate aldolase, and enzyme associated with anaerobic glycolysis pathways (Martínez-Fernández et al., 2008, 2010). While these enzymes are not directly associated with aerobic metabolism, they likely indicate a greater metabolic machinery required to maintain this capacity for activity, coupled with a greater aerobic requirement of the larger foot muscle. This increased metabolic capacity may come at the cost of heightened thermal sensitivity at the whole-organism level. Maintaining elevated metabolic rates at high temperatures is energetically costly, particularly under prolonged thermal stress, but may not be under strong selection in the mid- to lower-shore thermal environment occupied by the Wave ecotype.

As noted in our methodology, attrition rates of tethered snails in the field experiment were high. Although quantitative data on the causes of losses could not be collected, anecdotal observations suggest that losses resulted from a combination of ecological and non-ecological factors, with different relative importance across shore levels. Weaknesses in the superglue bond, resulting in snails escaping their tethers, was likely a source of attrition on the mid shore, where the attritional effect of wave action was high. Crab predation (Boulding et al., 2017) was a factor resulting in losses on the upper shore, evidenced by the presence of broken shell fragments attached to the tethers observed across both ecotypes, along with the presence of the crab *Pachygrapsus marmoratus* on the higher upper shore.

We did not find indications of heat-induced mortality, such as the presence of deceased but intact snails, or intact empty shells still attached to tethers. Thus, while thermal exposure on the upper shore was sufficient to induce physiological stress responses in the Wave ecotype within four days, it may not have been sufficient to cause mortality over this timeframe.

Longer-term studies have demonstrated reduced survival of Wave ecotype individuals transplanted to the upper shore (Boulding et al., 2017; Rolán-Alvarez et al., 1997), supporting the idea that prolonged exposure to upper-shore conditions imposes cumulative physiological and ecological constraints.

From an evolutionary perspective, our study provides support for the role for physiological divergence in limiting the ability of the Wave ecotype to colonise the upper shore, potentially acting as a pre-mating barrier and contributing to the maintenance of genetic differentiation between ecotypes (reviewed in Keller & Seehausen, 2012). However, as both ecotypes co-occur and hybridise within the mid shore, habitat incompatibility alone only partially explains maintaining ecotype divergence within this study system. Therefore, understanding the extent to which the two ecotypes physiologically differ within this (sympatric) hybrid-zone interface zone, vs. at different shore heights can therefore provide critical insights into other potential mechanisms by which thermal adaptation may influence ecotype divergence (Keller & Seehausen, 2012). A key feature of our experimental design was the inclusion of Crab ecotype snails from both mid- and upper-shore habitats, allowing us to separate the effects of shore height from those of ecotype identity. Previous work has provided mixed evidence regarding whether the Crab ecotype represents a single physiologically homogeneous population, or whether mid-shore Crab ecotype snails may be physiologically closer to those of the Wave ecotype (Rolán-Alvarez et al., 1997). Recent work (Raffini et al., 2025), conducted on the same populations used in our current study, has found that mid-shore Crab ecotype snails show greater genetic and morphological convergence with adjacent mid-shore Wave populations than do upper shore Crab ecotype snails, potentially either reflecting introgression across the hybrid interface, or local adaptation to mid-shore conditions. This led the authors of that study to call into question the identification of the Crab ecotype as a single homogenous biological entity across shore heights (Raffini et al., 2025). However, our data provides physiological support for the Crab ecotype as a single entity, as both Crab ecotype populations displayed similar cardiac profiles and neither displayed changes in cardiac activity following reciprocal transplant, contrasting with the Wave ecotype. Thus, our study supports a clear physiological distinction between the Galician Wave and Crab ecotypes, irrespective of shore height, and may suggest that physiological divergence is conserved in the mid-shore hybrid zone interface even in the face of potential gene flow and morphological convergence.

The eco-evolutionary mechanisms involved in maintaining this physiological divergence between ecotypes at the hybrid interface are unclear, particularly given the apparent equal ability of both ecotypes to survive on the mid-shore (Rolán-Alvarez et al., 1997). Potential explanations include genetic linkage between physiological, morphological, and behavioural traits. Chromosomal inversions play a key role in maintaining traits associated with ecotype divergence in *Littorina saxatilis*, including shell size and shell aperture size (Koch et al., 2021; Raffini et al., 2025). Inversions facilitate the persistence of adaptive genetic differences by suppressing recombination of affected genome areas during meiosis, meaning sets of co-adapted alleles are inherited as a single unit (Kirkpatrick, 2010). Chromosomal inversions could partly explain why physiological divergence appears to be more closely tied to ecotype identity than to shore height, for example if physiological differentiation is linked to the same inversions responsible for the distinct shell characteristics of the Galician Crab and Wave ecotypes. At broader geographic scales, chromosomal inversions are associated with repeated ecological divergence in *L. saxatilis* across both vertical shore-height gradients and wave exposure–predation gradients (Johannesson et al., 2024; Morales et al., 2019). However, the specific inversion sites associated with divergence across the two gradients are not necessarily the same (Johannesson et al., 2024; Morales et al., 2019), highlighting that more work is needed to directly link specific inversion sites to physiological traits (Koch et al., 2021), including in the Galician populations where both gradients occur in parallel and are synergistic (Raffini et al., 2025). Reduced physiological fitness of hybrids may also contribute to ecotype divergence at the hybrid interface (Keller & Seehausen, 2012), but physiological data on hybrids are extremely limited, with Rolán-Alvarez et al. (1997) reporting no clear survival differences between hybrids and parental lineages on the mid-shore. Future work focusing on hybrids could help clarify the role of physiological divergence in maintaining the ecotype system but would need to be paired with genetic work to establish the genetic heritage of individual hybrid animals, as this is extremely challenging based on visual inspection alone.

In conclusion, our results demonstrate that divergence in thermal physiology forms an important component of ecotype differentiation in Galician *Littorina saxatilis*, while also providing direct evidence of fitness consequences of physiological stress on the upper shore in the Wave ecotype, potentially restricting their vertical distribution across shore height and acting as an additional barrier to gene flow. Combined with previously documented differences in morphology, behaviour, and life-history traits, our findings suggest that adaptation to contrasting shore environments involves multiple trait complexes that are subject to selection in tandem. In this context, our findings highlight *L. saxatilis* as an example of ecological niche dimensionality and multidimensional divergent selection, whereby divergence and reproductive isolation is strengthened through the accumulation of selection across multiple traits and environmental dimensions (Chevin et al., 2014; Langerhans & Riesch, 2013; Nosil et al., 2009; Nosil & Sandoval, 2008; White & Butlin, 2021). Given the ubiquity of environmental gradients where multiple selective pressures covary across spatial scales in nature, comparisons of divergent ecotypes in *L. saxatilis* can offer insights into mechanisms of local adaptation and speciation which extend well beyond the intertidal to encompass a range of natural systems.

## Author attribution

CD, MT, ERA and JG conceived the experiment and obtained ASSEMBLE Plus funding. ERA and JG provided the thermal imaging camera, water bath, and drilling/ tethering equipment, and facilitated access to laboratory and field sites. MT and CD provided temperature loggers and heart rate monitoring equipment. JG, ERA, RL and CD installed field attachment tethers and temperature loggers. CD, RL and JG performed the field reciprocal transplant experiment and collected field heart rate and temperature data. CD and RL collected laboratory heart rate data. CD analyzed the data, prepared figures and tables, and wrote the original draft. All authors contributed to the manuscript and approved the final version for submission.

## Acknowledgements

We thank the Toralla Marine Station (ECIMAT-UVIGO) for access to and use of their facilities. We also thank Maria Huete, project manager and access officer at EMBRC-ES, for all the help with the ASSEMBLE Plus project that made this research possible.

## Data accessibility

Data and code associated with the manuscript will be made available at Zenodo upon publication.

## Appendix

**Table S1:**
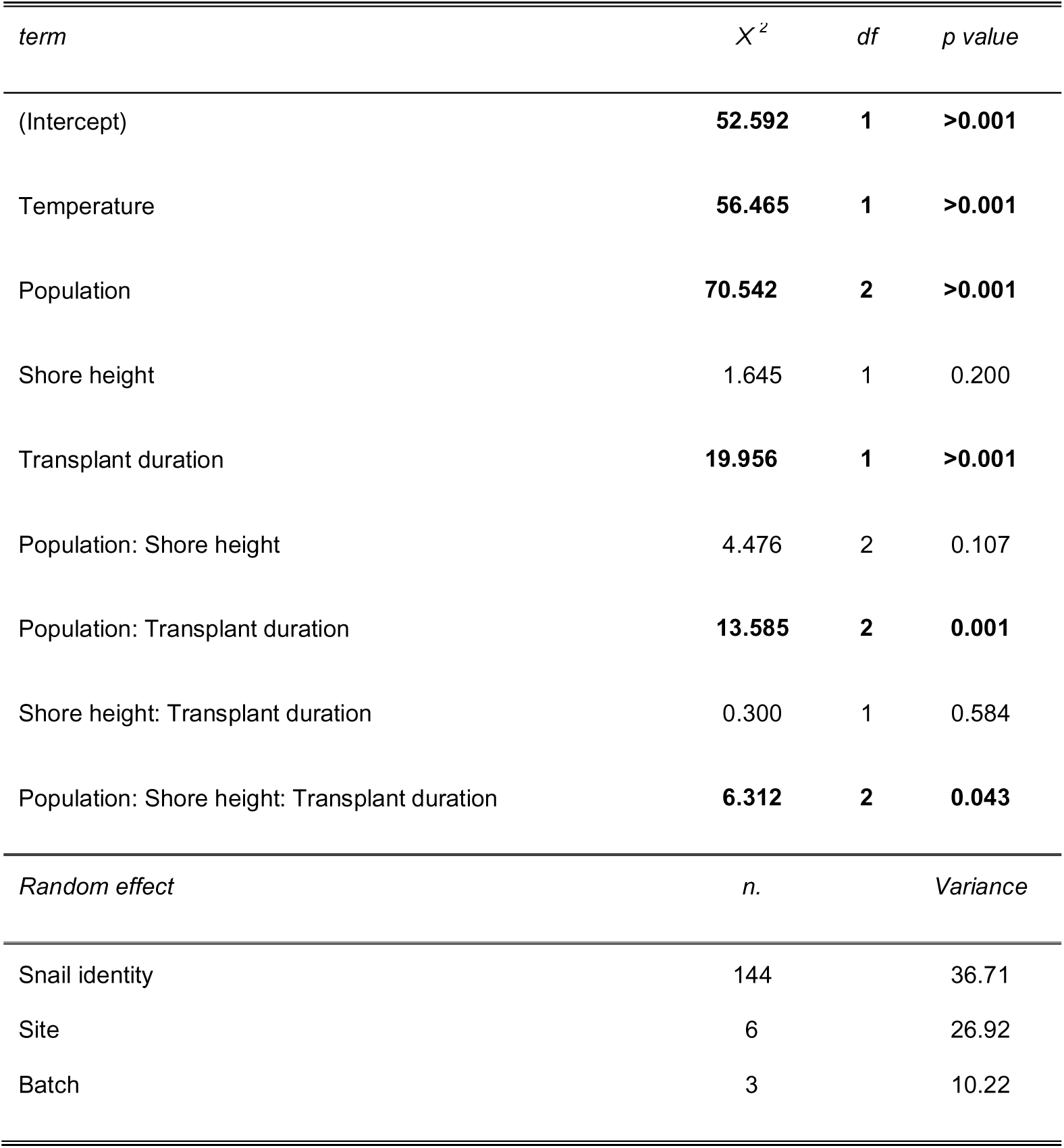
ANOVA table from statistical analysis of reciprocal transplant experiment.

**Table S2:**
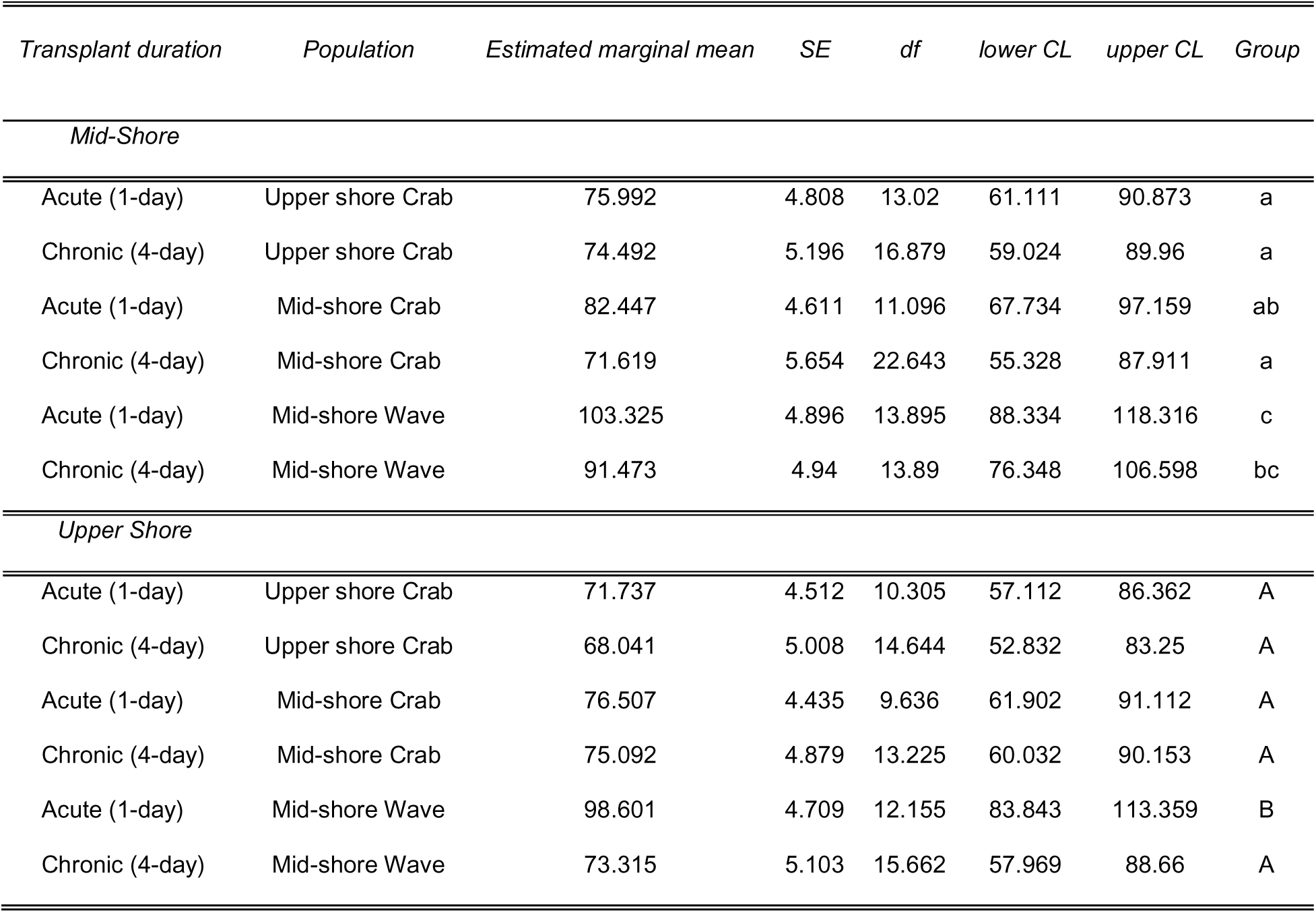
Pairwise contrasts from statistical analysis of reciprocal transplant experiment.

| *Transplant duration* | *Population* | *Estimated marginal mean* | *SE* | *df* | *lower CL* | *upper CL* | *Group* |
| --- | --- | --- | --- | --- | --- | --- | --- |
| *Mid-Shore* |  |  |  |  |  |  |  |
| Acute (1-day) | Upper shore Crab | 75.992 | 4.808 | 13.02 | 61.111 | 90.873 | a |
| Chronic (4-day) | Upper shore Crab | 74.492 | 5.196 | 16.879 | 59.024 | 89.96 | a |
| Acute (1-day) | Mid-shore Crab | 82.447 | 4.611 | 11.096 | 67.734 | 97.159 | ab |
| Chronic (4-day) | Mid-shore Crab | 71.619 | 5.654 | 22.643 | 55.328 | 87.911 | a |
| Acute (1-day) | Mid-shore Wave | 103.325 | 4.896 | 13.895 | 88.334 | 118.316 | c |
| Chronic (4-day) | Mid-shore Wave | 91.473 | 4.94 | 13.89 | 76.348 | 106.598 | bc |
| *Upper Shore* |  |  |  |  |  |  |  |
| Acute (1-day) | Upper shore Crab | 71.737 | 4.512 | 10.305 | 57.112 | 86.362 | A |
| Chronic (4-day) | Upper shore Crab | 68.041 | 5.008 | 14.644 | 52.832 | 83.25 | A |
| Acute (1-day) | Mid-shore Crab | 76.507 | 4.435 | 9.636 | 61.902 | 91.112 | A |
| Chronic (4-day) | Mid-shore Crab | 75.092 | 4.879 | 13.225 | 60.032 | 90.153 | A |
| Acute (1-day) | Mid-shore Wave | 98.601 | 4.709 | 12.155 | 83.843 | 113.359 | B |
| Chronic (4-day) | Mid-shore Wave | 73.315 | 5.103 | 15.662 | 57.969 | 88.66 | A |

